# 11β-hydroxysteroid dehydrogenase type 2 may mediate the stress-specific effects of cortisol on brain cell proliferation in adult zebrafish (*Danio rerio*)

**DOI:** 10.1101/2024.05.14.594160

**Authors:** E. Emma Flatt, Sarah L. Alderman

## Abstract

Stress-induced increases in cortisol can stimulate or inhibit brain cell proliferation, but the mechanisms behind these opposing effects are unknown. We tested the hypothesis that 11β-hydroxysteroid dehydrogenase type 2 (Hsd11b2), a glucocorticoid-inactivating enzyme expressed in neurogenic regions of the adult zebrafish brain, mitigates cortisol-induced changes to brain cell proliferation using one of three stress regimes: a single 1-min air exposure (acute stress), two air exposures spaced 24 h apart (repeat acute stress), or social subordination (chronic stress). Plasma cortisol was significantly elevated 15 min after air exposure and recovered within 24 h after acute and repeat stress, whereas subordinate fish exhibited significant and sustained elevations relative to dominant fish for 24 h. Following acute stress, brain *hsd11b2* transcript abundance was significantly lower 24 h after a single air exposure but was unchanged by repeat stress or social subordination. A sustained increase in brain Hsd11b2 protein levels occurred after acute stress, but not after repeat or chronic stress. Following acute and repeat stress, brain *pcna* transcript abundance exhibited a prolonged elevation, but was unaffected by social subordination. Interestingly, the number of telencephalic BrdU+ cells increased in fish after a single air exposure but was unchanged by repeat stress. Following acute and repeat stress, fish expressed lower brain *gr* and *mr* transcript abundance while subordinate fish exhibited no changes. Taken together, these results demonstrate stressor-specific regulation of Hsd11b2 in the zebrafish brain that could modulate rates of cortisol catabolism contributing to observed differences in brain cell proliferation.

**Summary Statement:** The steroid dehydrogenase, Hsd11b2, is dynamically regulated in the zebrafish brain during stress. This study provides novel evidence that variation in Hsd11b2 abundance may underscore stress-specific changes in forebrain neurogenesis.

## Introduction

Adult neurogenesis is the formation of new functional neurons from neural stem and progenitor cells (NSPCs) in the postnatal brain. Adult neurogenesis in the forebrain has been described in at least one species of each vertebrate class (Kaslin et al., 2008; Zupanc, 2021), however, considerable variation in the number of neurogenic niches and the rate of proliferation exists across taxa. Teleost fish have emerged as important model species in this field due to their comparatively greater capacity for lifelong neurogenesis (Zupanc, 2021). In the zebrafish (*Danio rerio*), for example, neurogenic niches are not restricted to the forebrain, but occur along the entire rostro-caudal axis of the brain (Zupanc et al., 2005) and the rate of NSPC proliferation far exceeds that observed in mammals (Hinsch and Zupanc, 2007). The functional significance of adult neurogenesis in teleost fish is an area of active study, but it is widely accepted to have both additive and reparative roles, i.e., supporting indeterminant growth (Zupanc and Horschke, 1995; Traniello et al., 2014) and injury repair (März et al., 2011; Lefevre et al., 2017), respectively. In addition, evidence that lifelong neurogenesis in the telencephalon contributes to learning and memory in fish (Ausas et al., 2019; Mazzitelli-Fuentes et al., 2022), as it does in mammals (van Praag et al., 1999; Lemaire et al., 2000; Hairston et al., 2005; Monteiro et al., 2014;), songbirds (Hall et al., 2014; Guitar and Sherry, 2018), and reptiles (LaDage et al., 2013), suggests that certain attributes of adult neurogenesis are conserved across vertebrate taxa.

Numerous endogenous factors can alter adult neurogenesis including the stress-induced release of glucocorticoid hormones. Glucocorticoids are pleiotropic steroid hormones that bind to both glucocorticoid and mineralocorticoid receptors (Gr and Mr) (Wendelaar Bonga, 1997; Mommsen et al. 1999; Aluru and Vijayan, 2009). Both Gr and Mr are widely expressed in the vertebrate brain (Wendelaar Bonga, 2011; Godoy et al., 2018), including in regions where NSPC proliferation occurs. Many studies in mammals suggest that acute transient increases in glucocorticoids stimulate brain cell proliferation (Thomas et al., 2006; Dagyte et al., 2009; Kirby et al., 2013; So et al., 2017), while chronically elevated glucocorticoids inhibit brain cell proliferation (Gould et al., 1998; Daygte et al., 2009). Research in fish supports a similar association between plasma cortisol, the principal glucocorticoid in fish, and brain cell proliferation. For example, the expression of *proliferating cell nuclear antigen* (*pcna;* a cell proliferation marker) was upregulated following brief air exposure in the whole brain of black rockfish (*Sebastes schlegelii*, Zhang et al., 2020) and after short-term confinement in the telencephalon of rainbow trout (*Oncorhynchus mykiss*, Johansen et al., 2012), concomitant with an increase in plasma cortisol. Conversely, the sustained increase in plasma cortisol resulting from a cortisol-laced diet or from social subordination reduced cell proliferation in the telencephalon (Sørensen et al. 2011; Sørensen et al., 2012; Tea et al. 2019). Nevertheless, findings that oppose these trends are not uncommon. For example, the transient increase in plasma cortisol following a series of acute stressors (10 min confinement, 1 min air exposure, and 5 min chase) was associated with fewer 5’-bromo-2’-deoxyuridine labelled cells (BrdU+, a thymidine analog) in the telencephalon of juvenile European sea bass (*Dicentrarchus labrax*) relative to unstressed controls (Fokos et al., 2017), whereas chronically increased plasma cortisol in brown ghost knifefish (*Apteronotus leptorhynchus*) coincided with greater numbers of BrdU+ cells in the periventricular zones compared with fish held in isolation (Dunlap et al., 2006). Importantly, pharmacological agents that inhibit cortisol synthesis or block Gr have successfully reversed the effects of stress on brain cell proliferation in fish (Dunlap et al., 2011; Tea et al., 2019). Altogether, the results of these studies support a role for cortisol in regulating neurogenesis along a continuum from stimulatory to inhibitory, however, the mechanisms that underpin this variable regulation are unclear. Intracellular glucocorticoids can be catabolized by the enzyme 11β-hydroxysteroid dehydrogenase type 2 (Hsd11b2), which converts cortisol (humans, fish) or corticosterone (rodents, birds) to their inert forms, cortisone or 11-dehydrocorticosterone, respectively (Chapman et al., 2013). Cortisone is then converted to 20β-hydroxycortisone by 20β-hydroxysteroid dehydrogenase type 2 (Hsd20b2) for excretion (Tokarz et al., 2012, 2013). In addition, the enzyme Hsd11b2 also demonstrates other functions in steroid hormone pathways, including generating the principal fish androgen, 11-ketotestosterone (Tsachaki et al., 2017). At the cellular level, the activity of Hsd11b2 may account for variable responses to cortisol signaling by buffering intracellular cortisol levels and the downstream activation of Gr and Mr.

Indeed, previous work in mammals has shown that HSD11B2 in the placenta and in the fetal brain are critical for protecting brain development during maternal stress by preventing glucocorticoid-induced inhibition of neurogenesis (Diaz et al., 1998; Wyrwoll et al., 2015). Intriguingly, Hsd11b2 is broadly expressed in the brains of adult zebrafish (Alderman and Vijayan, 2012), including in regions of significant neurogenic activity in the telencephalon (Zupanc, 2006), and acute stress increased Hsd11b2 enzymatic activity in the brain for at least 24 h post-stress (Alderman and Vijayan, 2012). Nevertheless, a role for Hsd11b2 in regulating fish neurogenesis during stress is unexplored. Therefore, this study tested the hypothesis that variation in brain Hsd11b2 underpins the stressor-specific effects of cortisol on NSPC proliferation in the zebrafish brain.

## Materials and Methods

### Animals

Adult wild-type zebrafish (*Danio rerio*) were obtained from a local supplier and held at the Hagen Aqualab (University of Guelph, Ontario, Canada). Fish were maintained at 27°C with a 13L:11D photoperiod and fed twice daily with GEMMA Micro 300 (ZEBCARE B.V., Nederweert, The Netherlands). The use and care of these animals were approved by the University of Guelph’s Animal Care Committee in accordance with guidelines of the Canadian Council for Animal Care.

### Experiment 1: Stress-specific regulation of hsd11b2 in the brain

Fish were exposed to one of three stress regimes: a single 1-min air exposure (acute stress), two 1-min air exposures spaced 24 h apart (repeat acute stress), or social subordination (chronic stress). For acute stress and repeat acute stress, a total of 91 adult mixed sex zebrafish were randomly distributed across fourteen 2 L tanks (N=6-7 per tank; 2 tanks per time point). Following a 1-week acclimation, fish were either immediately euthanized as described below (time 0) or subjected to a standardized 1-min air exposure by simultaneously transferring all fish in a tank to a fish net and holding them in the air for exactly 1 min, as previously described (Alderman and Vijayan, 2012). The fish were then returned to the tank to recover for 15 min, 6 h, 24 h, or 48 h. The fish from four experimental tanks underwent a second 1-min air exposure at 24 h followed by recovery for 15 min or 24 h (repeat acute stress) to determine whether any changes in Hsd11b2 expression would be augmented or suppressed by a second stressor. At the determined recovery time, all fish in a tank were rapidly and simultaneously killed using an overdose of ice-cold buffered tricaine methane sulfonate (MS-222; 0.3 g L^-1^ MS-222, 0.6 g L^-1^ NaHCO_3_). The tail was severed, and blood collected by gravity flow as described by Babaei et al. (2013), followed by blood centrifugation to isolate plasma (N = 13 per timepoint). The brains were then removed, snap frozen on dry ice, and stored at −80°C for future analysis of gene (N=8 per timepoint) or protein expression (N= 5 respectively per timepoint). Note that for acute and repeat stress gene expression, the telencephalons were isolated from the remaining brain regions (2 telencephalons/tube, N = 4) to examine region-specific changes.

For chronic stress, a total of 24 adult male zebrafish were anesthetized using MS-222 (0.15 g L^-1^ MS-222, 0.3 g L^-1^ NaHCO_3_) and fin-clipped for identification. Fish were size-matched based on weight (mean dominant = 0.80 ± 0.02 g, mean subordinate = 0.81 ± 0.02 g, mean difference between pairs = 0.006 ± 0.003 g) and fork length (mean dominant = 4.22 ± 0.04 cm, mean subordinate = 4.27 ± 0.04 cm, mean difference between pairs = 0.04 ± 0.20 cm). Each pair of fish was acclimated for 24 h in a 2.6 L tank, with one individual on either side of a perforated opaque divider. Each experimental tank contained an air stone and a constant inflow of filtered well water at ∼ 27°C supplied on both sides of the divider. The barrier was then removed, and the fish were allowed to interact for 5 min to establish the dominant-subordinate hierarchy. The hierarchy was considered to have formed when one fish started to perform retreats and ceased using aggressive behaviours. The behaviour of each fish was then recorded using an established scoring system (Filby et al., 2010; Paull et al., 2010; Jeffrey and Gilmour, 2016; Tea et al., 2019). Briefly, behavioural scores were taken twice daily where acts of aggression (biting, chasing) and number of retreats were tallied for each fish in a pair during a 5 min observation period. The positions of the fish in the tank and whether it fed first or second were noted. The fish were allowed to interact for 24 h and then were killed and tissues collected as above. This timing was chosen based on Tea et al. (2019) where reduced forebrain cell proliferation was observed in subordinate fish injected with BrdU after 24 h of social interaction, and to permit comparison with the 24 h recovery timepoint of the acute and repeat acute stress experiment. A separate cohort of fish was directly sampled from group housing to serve as a control group with presumed neutral social status (N = 13).

### Experiment 2: Effect of acute and repeat acute stress on cell proliferation in the telencephalon

A total of 12 adult mixed sex zebrafish were randomly distributed across three 2L tanks (N = 4 per tank) and acclimated for 1 week. Following acclimation, fish in an experimental tank were either netted and immediately anesthetised (time 0) or subjected to a single or repeated 1-min air exposure (see *Experiment 1*) before undergoing light anesthesia by gradual cooling to 12°C. Each fish was given a single intraperitoneal injection of BrdU (Sigma-Aldrich, Saint Louis, MO, USA; 40 µL/g body weight) and then allowed to recover for 1.5 h (timing informed by results of *Experiment 1*). The fish were killed as above and the brains prepared for cryosectioning exactly as previously described (Tea et al., 2019).

### Plasma cortisol

Plasma was thawed on ice, diluted 30X with 1X assay buffer and used in duplicate reactions to quantify cortisol concentration using a commercially available enzyme-linked immunosorbent assay (ELISA) (Neogen, Lexington, KY, USA) exactly according to manufacturer instructions. Samples that fell outside the range of the standard curve could not be re-assayed due to limited plasma availability, and were therefore removed from further analysis. Intra-assay variation was 3.3% across all plates. Inter-assay variation was 9.5% for acute stress samples; all chronic stress samples were assayed on a single plate.

### Western blot: quantifying Hsd11b2 protein abundance

Total protein was extracted from whole brain samples by homogenizing in 100 µL of 1X RIPA buffer containing protease inhibitors (cOmplete Mini Protease Inhibitor Cocktail, Hoffmann La Roche Limited, Mississauga, ON, Canada) and prepared for Western blotting exactly as previously described (Dindia et al., 2017). For each sample, 20 µg total protein was loaded on a 10% SDS-polyacrylamide gel, separated by SDS-PAGE and transferred to 0.45 µm nitrocellulose membrane (Bio-Rad Laboratories Ltd., Montreal, QC, Canada; 30V x 16 h). Each gel contained a loading control (pooled brain homogenate) and all representative timepoints (acute, repeat) or social statuses (chronic) for a given experiment. Following transfer, the membranes were blocked in 1X fish gelatin blocking buffer (Biotium Inc., CA, USA) for 1 h at room temperature and then incubated overnight at 4°C in chicken-anti-Hsd11b2 (Genetel, Madison, WI, USA) diluted to 0.5 µg mL^-1^ in 2% skim milk prepared in Tris buffered saline containing 0.05% Triton X-100 (TBST). The primary antibody was custom designed to recognize a conserved region of rainbow trout and zebrafish Hsd11b2 (Best et al., 2023), and was a generous gift from Dr. K.M. Gilmour (University of Ottawa, Canada). The antibody was fully validated for specificity and linear quantitation prior to use in these experiments (Fig. S1). Immunodetection was completed with a 1 h incubation in HRP-conjugated secondary antibody (goat-anti-chicken IgY H&L (HRP); cat. no. ab97135, Abcam, Cambridge, UK) diluted to 0.13 µg mL^-1^ in 2% skim milk in TBST, followed by chemiluminescent detection using Clarity Western Enhanced Chemiluminescence Substrate (Bio-Rad) on a ChemiDoc MP Imaging System (Universal Hood III, Bio-Rad). The membranes were then stripped (2 mM glycine, 0.1% sodium dodecyl sulphate, 1% Tween 20; pH 2.2) for 10 min at room temperature and washed (2 x 10 min in phosphate buffered saline, PBS), 1 x 5 min in TBST, 1 x 5 min in TBS) prior to total protein staining using SYPRO Ruby Protein Blot Stain (Invitrogen Canada Inc, Burlington, ON, Canada) according to the manufacturer’s instructions. The protein abundance of each sample was quantified as band intensity (arbitrary units, AU) using the Image Lab Software Version 6.1 (Bio-Rad) and normalized to total protein.

### Quantitative RT-qPCR

Total RNA was extracted from frozen brain tissue using TRIzol Reagent (Invitrogen) following manufacturer instructions. The purity and concentration were verified (NanoDrop 2000 Spectrophotometer; Thermo Fisher Scientific, Waltham, MAS, USA) and then 1 µg of total RNA was treated with DNase I prior to cDNA synthesis using the High-Capacity cDNA Reverse Transcription kit (Applied Biosystems, CA, USA), all according to manufacturers’ instructions. To test the efficacy of the DNase treatment, non-reversed transcribed (non-RT) controls were included for a subset of random samples (∼10%) by omitting the MultiScribe RT enzyme in the cDNA synthesis reaction. Quantitative reverse transcription polymerase chain reaction (RT-qPCR) was used to quantify the gene expression of *elongation factor 1α (ef1α)*, *ribosomal protein L8 (rpl8)*, *hsd11b2*, *pcna*, *gr,* and *mr* separately in duplicate 15 µL reactions containing 1x Sso Advanced Universal SYBR Green Supermix (Bio-Rad), gene specific primer pair (Table 1) and 5 µL of template (diluted cDNA, non-RT, or water). Cycling conditions were as specified by the manufacturer and included a final melt curve. Mean cycle threshold (Ct) values were fitted to the antilog of standard curves generated for each primer pair using serially diluted cDNA. The transcript abundance of each gene of interest was then normalized to the mean expression of two housekeeping genes, *ef1α* and *rpl8*, which were stably expressed across all treatments (one-way analysis of variance, ANOVA).

**Table 1.**
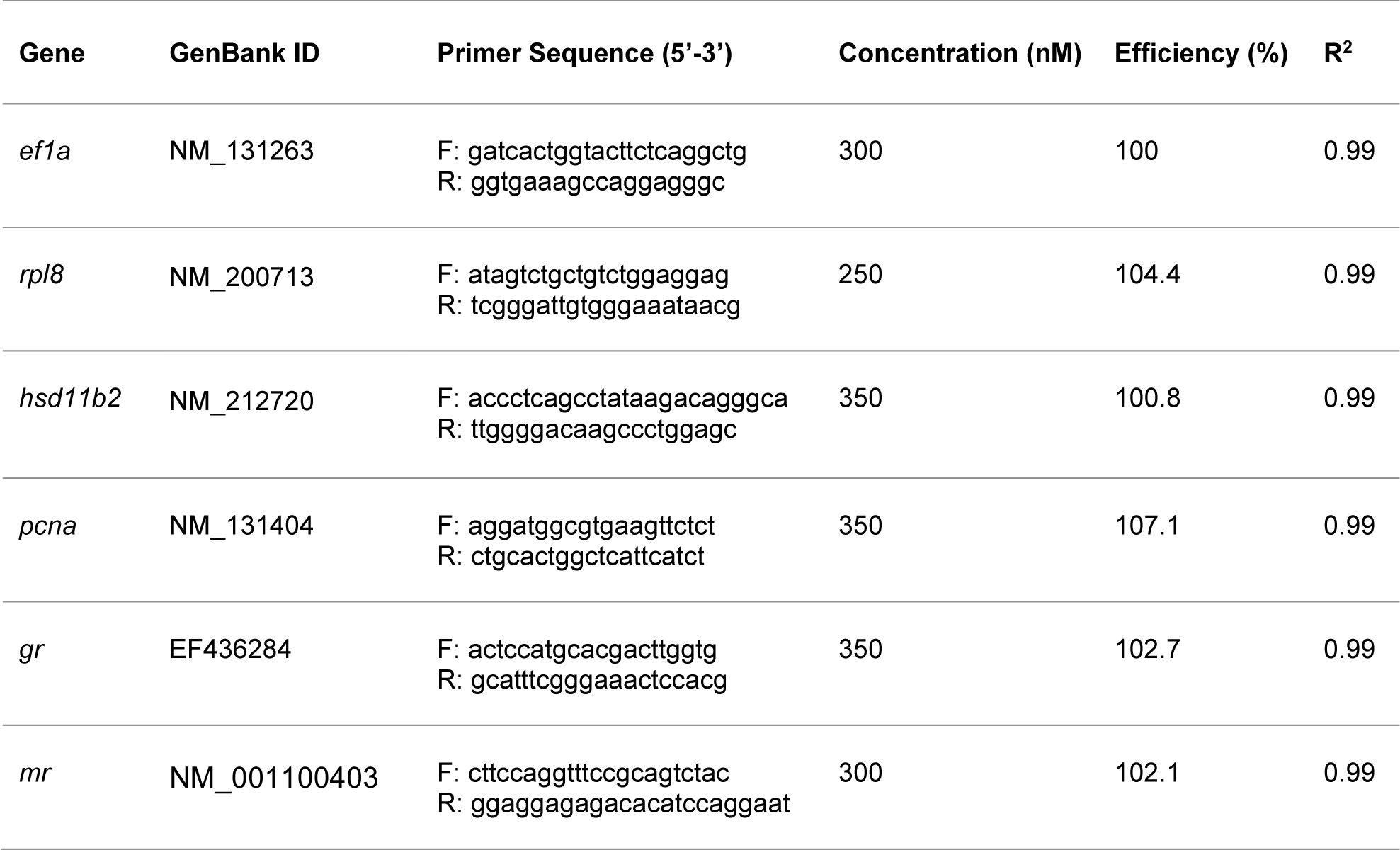
Gene-specific forward (F) and reverse (R) primer sequences that were used for RT-qPCR analysis. *ef1a*, elongation factor 1-alpha; *rpl8*, ribosomal protein L8; *hsd11b2*, 11β-hydroxysteroid dehydrogenase type II; *pcna*, proliferating cell nuclear antigen; *gr*, glucocorticoid receptor; *mr*, mineralocorticoid receptor.

### Immunohistochemistry

Slides were thawed at room temperature and then rehydrated in PBS prior to antigen retrieval in 10 mmol L^-1^ sodium citrate buffer (pH = 6) with 0.05% Tween-20 at 65°C for 30 min. Slides were cooled to room temperature for 15 min, then immersed in 2 mol L^-1^ HCL for 15 min followed by 2 x 5 min washes in 0.1 mol L^-1^ borate buffer (pH = 8.5). The sections were permeabilized for 3 x 5 min in PBS containing 0.05% Triton X-100 (PBST), and then blocked with 5% goat serum (Invitrogen) in PBST for 45 min at room temperature. BrdU-labelled cells were visualized using mouse-anti-BrdU (cat. no. G3G4 (anti-BrdU), Developmental Studies Hybridoma Bank, Iowa City, IA, USA) diluted to 1:100 in blocking buffer (overnight, 4°C) and Alex Fluro 488-conjugated goat-anti-mouse IgG (cat. no. 115-545-003, Jackson Immuno Research, West Grove, PA, USA) for 3 h at room temperature in the dark. Every third section through the telencephalon (12 sections/fish) was imaged on a Nikon ECLIPSE Ti2 (Nikon Instruments Inc., New York, USA). All BrdU+ cells were tallied and normalized to cross-sectional area (µm^2^), which was stable across all experimental groups (one-way ANOVA).

### Statistical Analysis

Social rank was assigned based on the behavioural scores of each fish using a principal components analysis (PCA), low behaviour scores were associated with dominance (PC1 eigenvalue = 1.7). Therefore, within each pair, the fish with the lower behaviour score was assigned dominant status (mean behaviour score = −1.65 ± 0.05) and the fish with the higher score was assigned subordinate status (mean behaviour score = 1.65 ± 0.06).

Differences in plasma cortisol, normalized mRNA abundance (*hsd11b2, pcna, gr, mr)*, and Hsd11b2 protein expression between recovery timepoints (acute stress, repeat stress) or between social status categories (chronic stress), as well as brain cell proliferation (BrdU+ cells/µm^2^) were determined using one-way ANOVAs followed by Tukey’s post-hoc tests. Data that did not meet the assumptions of normality (Shapiro-Wilk test) or equal variance (Bartlett test) were either log transformed or analyzed using non-parametric tests (Kruskal-Wallis followed by Dunn’s post-hoc test). Due to low RNA yield in telencephalon samples, *gr* and *mr* mRNA abundance could not be quantified at 6 h recovery from acute stress. Any datapoints greater than or less than the 1.5 x interquartile range from the upper or lower quartile, respectively, were removed prior to statistical analysis (no more than two outliers were removed from any treatment group). For transparency, all outliers are indicated on graphs as black triangle datapoints. All statistical analyses were performed in R Statistical Software (v4.2.2; R Core Team 2022) and α was set to 0.05. Figures were produced using GraphPad Prism version 10.2.0 (GraphPad Software, Boston, Massachusetts. USA).

## Results

### The effects of stressor exposure on plasma cortisol

Plasma cortisol levels changed over time after zebrafish were exposed to a 1-min air exposure stressor (F_4,51_ = 20.17, *P* < 0.0001; Fig. 1A). At 15 min post-stress, fish experienced a 75-fold increase in plasma cortisol levels relative to baseline (*P* < 0.0001). At 6 h post-stress, plasma cortisol levels remained elevated relative to baseline (*P* < 0.0001) but returned to pre-stress levels by 24 h (*P* > 0.5). Similarly, repeat stress exposure induced a 48-fold increase in plasma cortisol levels at 15 min recovery relative to baseline (H = 25.297, *P* < 0.0001, d.f. = 2; Fig. 1B), with complete recovery to baseline by 24 h (*P* = 0.0002).

**Figure 1.**
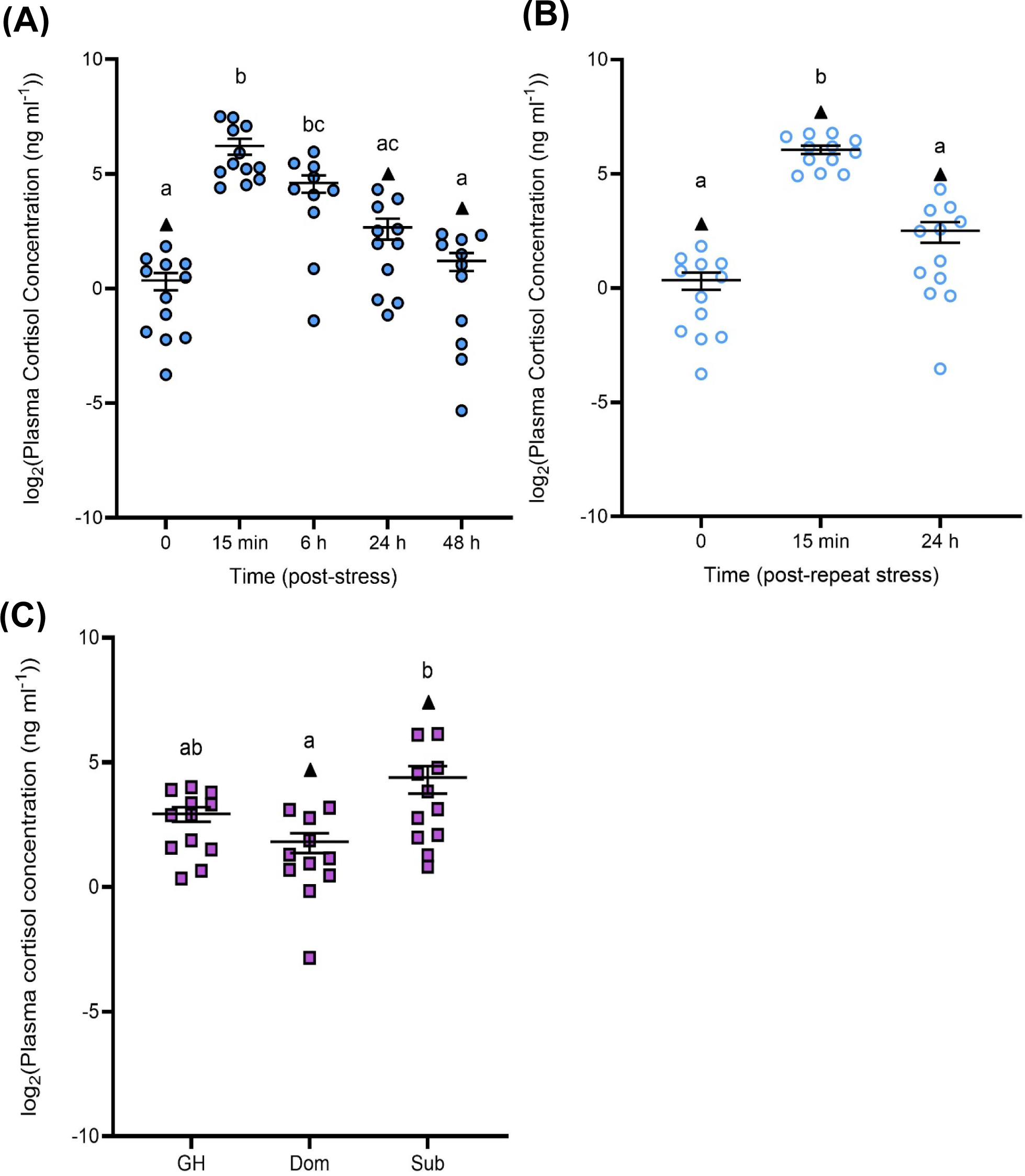
The effects of stress exposure on plasma cortisol. (ng ml^-1^)**. (A)** Zebrafish were exposed to air for 1-min and allowed to recover for up to 48 h (filled blue circles). (B) For a separate group of fish, the stressor was repeated 24 h after the first stressor and fish were allowed to recover for 15 min or 24 h (open blue circles). **(C)** Differences in the plasma cortisol of dominant (Dom) and subordinate (Sub) zebrafish after 24 h of social interaction; group-housed (GH) fish are included to act as a control (closed purple squares). Datapoints identified as statistical outliers were removed prior to analysis, and are shown as black triangles. Each symbol represents an individual fish, with a solid line and whiskers representing the mean ± SEM (acute stressor n = 10-13; repeat stressor n =12; social stress n = 11-13). Differences in plasma cortisol levels following stressor exposure were determined either using a one-way ANOVA and Tukey’s *post-hoc* test, or Welch’s ANOVA with a Kruskal-Wallis or Dunn’s *post-hoc* test (*P* < 0.05). Letters denote significant differences among groups within a panel; groups sharing the same letter are not significantly different from one another.

Plasma cortisol levels were affected by social status (F_2, 31_ = 5.528, *P* = 0.0090; Fig 1C). Subordinate fish experienced a 6.3-fold increase in plasma cortisol levels relative to dominant fish (*P* = 0.0073). Plasma cortisol of group-housed fish was intermediate between dominant and subordinate fish (*P* > 0.05).

### The effects of stress and recovery on hsd11b2 expression in the brain

Temporal changes in brain *hsd11b2* mRNA abundance (F_4, 30_ = 8.952, *P* < 0.0001, Fig. 2A) and Hsd11b2 protein expression (H = 11.403, *P* = 0.0224, d.f. = 4; Fig. 2B) were observed following 1-min air exposure stress. The mRNA abundance of *hsd11b2* was unchanged at 15 min and 6 h recovery (*P* > 0.05). After 24 h and 48 h recovery, *hsd11b2* mRNA abundance was 2-fold lower than 15 min (*P* = 0.0004, and *P* = 0.0036) and 6h post-stress (*P* = 0.0012, and *P* = 0.0110). Similar trends in *hsd11b2* mRNA abundance were found in the pooled telencephalons, although it did not reach statistical significance (F_4,12_ = 2.63, *P* = 0.0870; Fig. S3A). Brain Hsd11b2 protein abundance was 1.7-fold higher than pre-stress levels by 6 h recovery (*P* =0.0142) and remained elevated through to 48 h post-stress (*P* = 0.0051). In contrast, repeat stress exposure did not significantly affect brain or telencephalon *hsd11b2* mRNA abundance (F_2, 20_ = 2.192, *P* = 0.1378; F_2, 7_ = 2.107, *P* = 0.1920; Fig. 2C, S3B, respectively); however, Hsd11b2 protein abundance was affected (F_2,10_ = 15.12, *P* = 0.0009; Fig. 2D). Specifically, Hsd11b2 was 2.3-fold higher than baseline at 15 min recovery (*P* = 0.0008) but returned to pre-stress levels by 24 h recovery.

**Figure 2.**
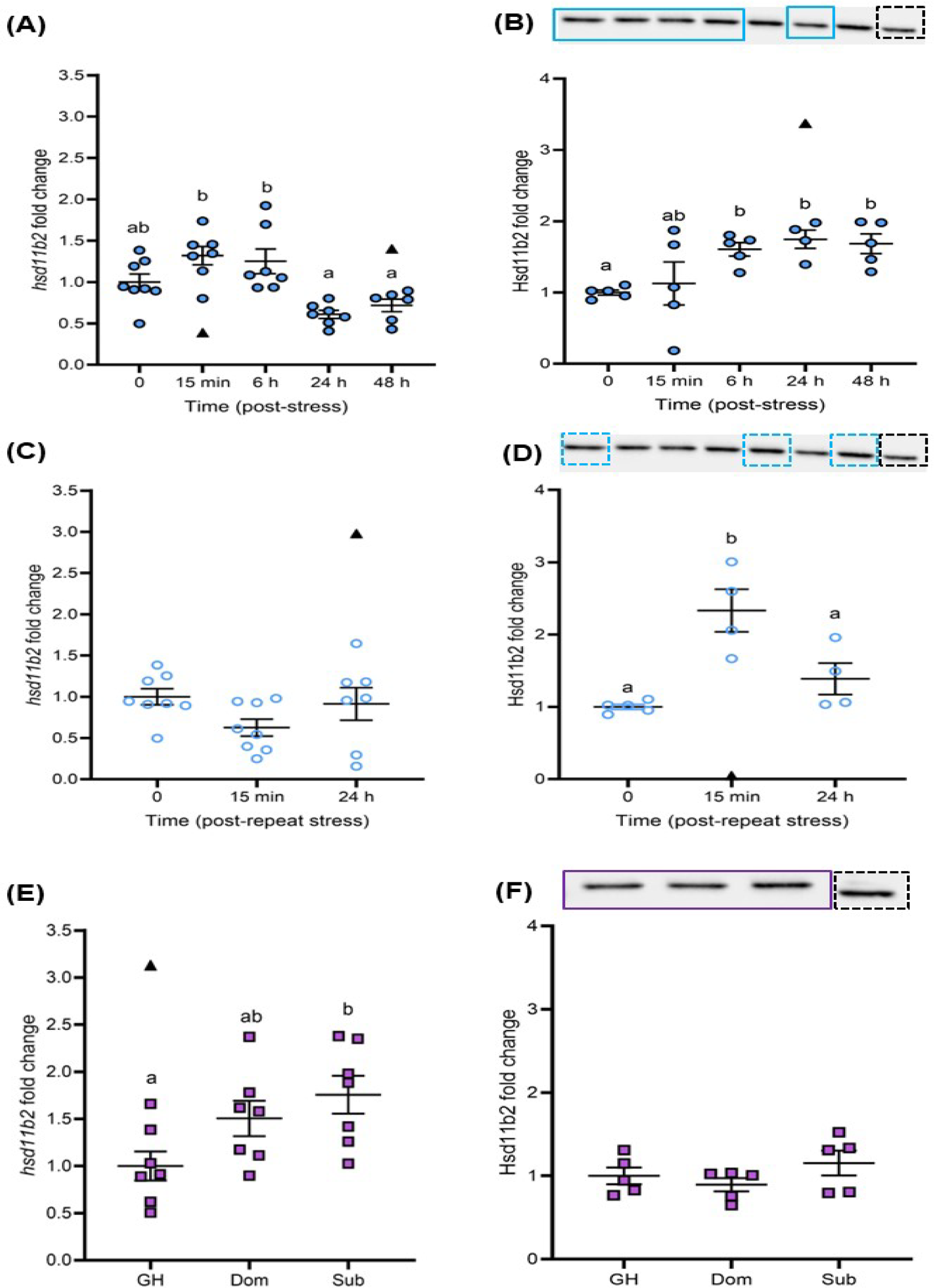
The effects of stress exposure on *hsd11b2* mRNA and Hsd11b2 protein abundance in the adult zebrafish brain. Brain *hsd11b2* transcript and Hsd11b2 protein abundance following exposure an acute stressor **(A, B)**, repeat acute stressor **(C, D)**, and chronic stressor **(E, F)**. A representative western blot is shown for all protein data, with the internal control marked by a dashed black box. In **(B)**, bands inside the blue boxes coincide with 0, 15 min, 6 h, 24 h, and 48 h post-stress, respectively. In **(D)**, bands inside the dashed blue boxes coincide with 0, 15 min, and 24 h post-repeat stress, respectively. In **(F)**, bands inside the purple box coincide with group housed (GH), dominant (Dom), and subordinate (Sub) fish, respectively. Transcript abundances were normalized to the mean expression of the two housekeeping genes (*ef1α* and *rpl8*). Protein expression of Hsd11b2 was normalized to total protein. Data is shown as individual data points with a solid line and whiskers representing the mean ± SEM (*hsd11b2*, acute stress n = 6-8, repeat stress n =7-8, social stress n = 7; Hsd11b2, acute stress n=4-5, repeat stress n=4-5, social stress n = 5). Datapoints identified as statistical outliers were removed prior to analysis, and are shown as black triangles. Differences in *hsd11b2* mRNA and Hsd11b2 protein levels following stressor exposure were determined either using a one-way or Welch’s ANOVA followed by a Tukey’s *post-hoc* test, or a Kruskal-Wallis followed by a Dunn’s *post-hoc* test (*P* < 0.05). Letters indicate significant differences among groups in each panel; groups which have the same letter are not significantly different from one another.

Social status had differential effects on brain *hsd11b2* mRNA abundance (F_2, 18_ = 4.48, *P* = 0.0264; Fig. 2E). Although *hsd11b2* mRNA abundance was similar between dominant and subordinate fish, it was 1.8-fold higher in subordinate fish relative to group-housed (*P* = 0.0260). There were no significant differences in brain Hsd11b2 protein abundance between dominant, subordinate, or group-housed fish (F_2, 12_ = 1.321, *P* = 0.3030; Fig. 2F).

### The effects of stress and recovery on brain cell proliferation

Brain *pcna* mRNA abundance was significantly different among timepoints following acute stress exposure (*H* = 23.346, d.f. = 4, *P* = 0.0001; Fig. 3A). The abundance of *pcna* was 3-fold higher than pre-stress levels after 15 min recovery (*P* < 0.0001) and remained elevated through 24 h recovery (*P* = 0.0199) but returned to baseline by 48 h recovery. Similarly, telencephalon *pcna* mRNA abundance was significantly affected by acute stress exposure (H =9.691, *P* = 0.0460, d.f. = 4; Fig. S4A). The *pcna* mRNA abundance of the pooled telencephalons showed a 5-fold increase at 15 min post-stress (*P* = 0.0019) but returned to baseline levels by 6 h recovery. Repeat stress also affected brain *pcna* mRNA abundance (F_2, 18_ = 27.78, *P* < 0.0001; Fig. 3B), where *pcna* mRNA levels were at least 1.7-fold higher than baseline levels at 15 min and 24 h recovery (*P* < 0.0001). A similar trend was found in pooled telencephalons, where *pcna* mRNA abundance was 2-fold higher at 15 min and 24 h post-repeat stress relative to baseline but did not reach statistical significance (*H* = 5.6, *P* = 0.0608, d.f. = 2; Fig. S4B). Social status did not affect brain *pcna* mRNA abundance (F_2,17_ = 0.943, *P* = 0.4090; Fig. 3C).

**Figure 3.**
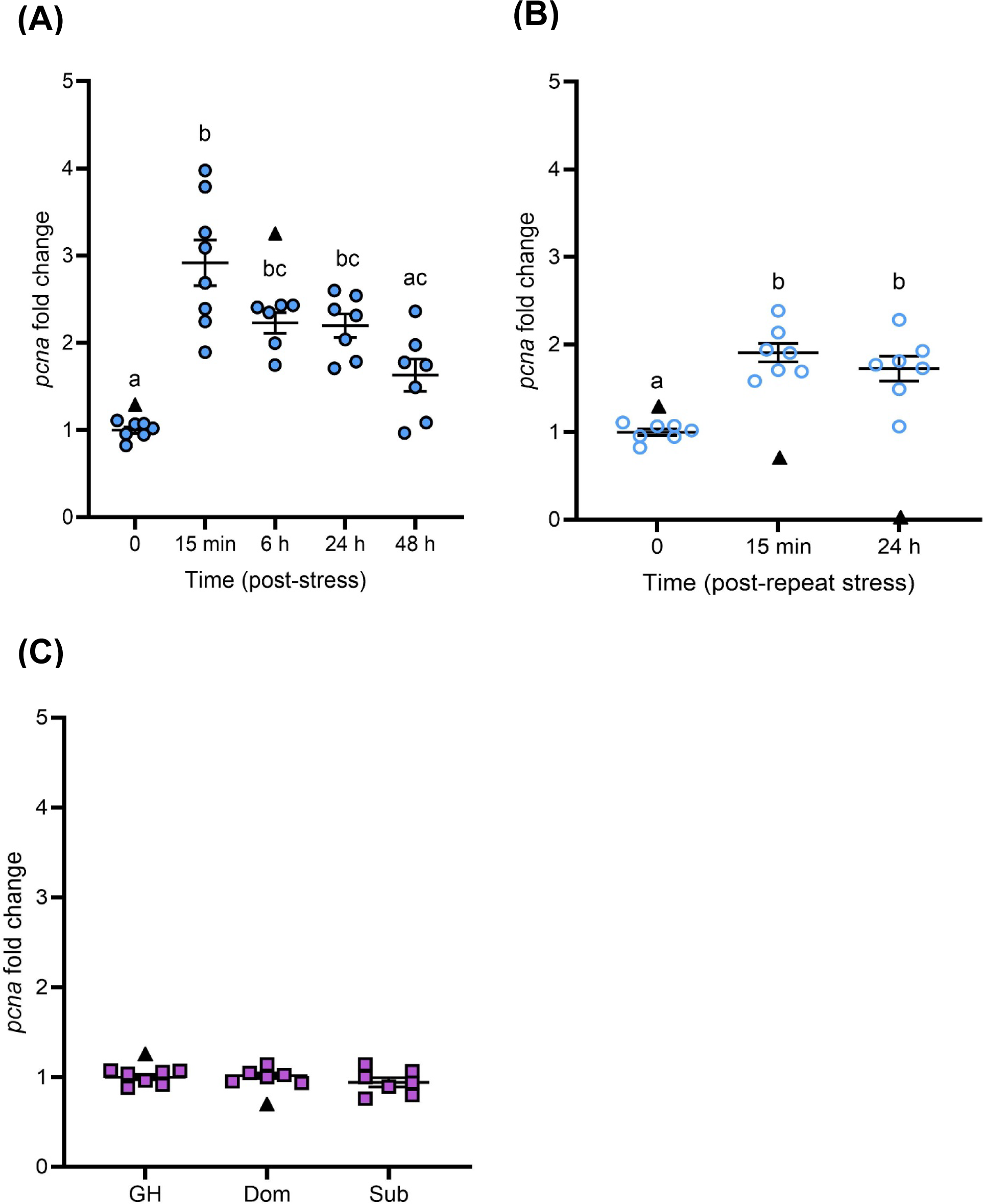
Effects of stress exposure on *pcna* transcript abundance in the adult zebrafish brain. **(A)** Zebrafish were exposed to air for 1-min and allowed to recover for up to 48 h (closed blue circles). **(B)** For a separate group of fish, the stressor was repeated 24 h after the first stressor and fish were allowed to recover for 15 min or 24 h (open blue circles). **(C)** Dominant (Dom) and subordinate (Sub) zebrafish after 24 h of social stress; group-housed (GH) fish are included as a control (purple squares). Datapoints identified as statistical outliers were removed prior to analysis, and are shown as black triangles. Gene transcript abundance was normalized to the mean expression of the two housekeeping genes (*ef1α* and *rpl8*). Data is shown as individual data points with a solid line and whiskers representing the mean ± SEM (*pcna* transcript abundance, acute stress n = 6-8; repeat stress n = 7; social stress n = 6-7). Changes in *pcna* transcript abundance following stress exposure were determined using a one-way ANOVA with a Tukey’s *post-hoc* test or a Kruskal-Wallis with a Dunn’s *post-hoc* test (acute and repeat stress *P* < 0.05; social stress *P* > 0.05). Letters denote significant differences among groups within a panel; groups which have the same letter are not significantly different from one another.

The number of BrdU+ cells in the telencephalon changed following an air exposure stressor (F_2,7_ = 6.215, *P* = 0.0281; Fig. 4). Specifically, the number of BrdU+ cells in the telencephalon was 1.6-fold greater at 1.5 h recovery from the acute stress relative to unstressed controls (*P* = 0.0300). Following 1.5 h recovery from the repeat stressor, the number of BrdU+ cells in the telencephalon was intermediate between the acute stress fish and unstressed controls (*P* = 0.1300, *P* = 0.2700).

**Figure 4.**
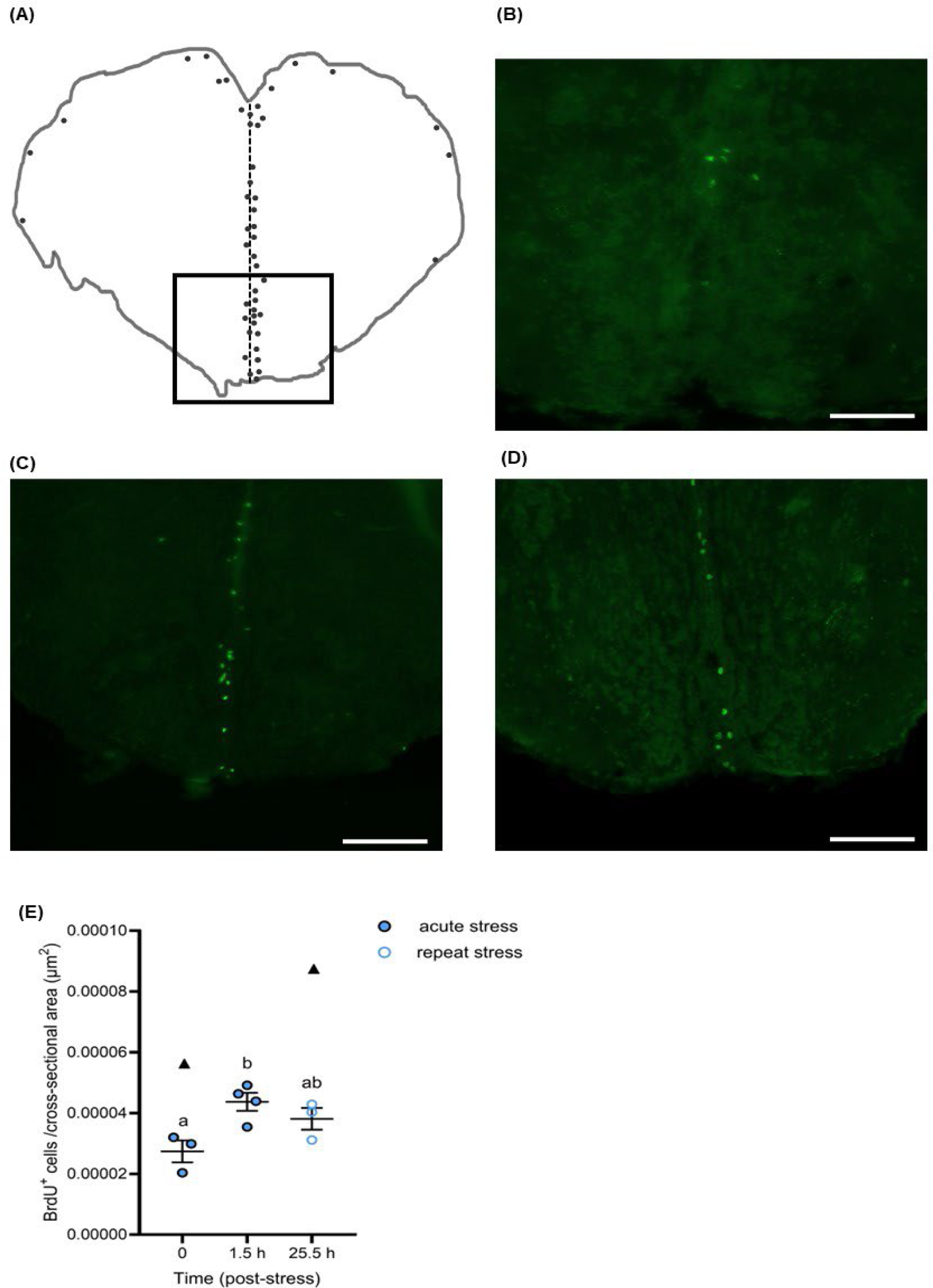
The effects of a single and repeat stressor on brain cell proliferation in the telencephalon of adult zebrafish brain. **(A)** Line drawing traced from original images of cross-sections following immunohistochemical detection, where the vertical dashed line represents the midline of the section and grey dots represent common locations of BrdU+ cells. The approximate corresponding level in the zebrafish brain atlas is the rostral telencephalon Level 71 (Wullimann et al., 1996). The boxed region indicates the area of the representative photomicrographs of cross-sections through the zebrafish telencephalon of **(B)** time 0 (baseline), **(C)** 1.5 h recovery following a single stressor, and **(D)** 1.5 h recovery from a repeat 1-min air exposure stressor 24 h after the initial stressor (i.e., 25.5 h from time 0); scale bars =100 µm. The number of BrdU+ cells (green) in each serial cross-section for individual fish were tallied and normalized to cross-sectional area (µm^2^), and then used to estimate cell proliferation **(E)**. Data is shown as individual data points with a solid line and whiskers representing the mean ± SEM (n = 3-4). Datapoints identified as statistical outliers were removed prior to analysis, and are shown as black triangles. Changes in cell proliferation following stress exposure were determined using a one-way ANOVA with a Tukey’s *post-hoc* test (*P* < 0.05). Letters denote significant differences among timepoints; timepoints which have the same letter are not significantly different from one another.

### The effects of stress and recovery on the transcript abundance of gr and mr in the brain

Brain *gr* transcript abundance was significantly different among timepoints following acute stress exposure (F_4,29_ = 10.02, *P* < 0.0001; Fig. 5A). The *gr* mRNA abundance was 1.4-fold lower than pre-stress levels as early as 15 min recovery, however this decrease did not reach statistical significance until 6 h, 24 h and 48 h (∼1.7-fold, P =0.0169, P = 0.0006, and *P* < 0.0001, respectively). The transcript abundance of *gr* in the pooled telencephalons was significantly affected by acute stress (*H* = 9.4762, *P* = 0.0236, d.f. = 3; Fig. S5A) where a significant decrease in *gr* mRNA abundance was observed at 48 h post-stress relative to baseline (*P* = 0.0046). Repeat stress significantly affected brain *gr* mRNA abundance (F_2, 19_ = 5.524, *P* = 0.0128; Fig. 5B), where a significant 1.5-fold decrease in *gr* mRNA abundance occurred at 24 h recovery relative to baseline (*P* = 0.0140). In contrast, *gr* mRNA abundance within the pooled telencephalons was not significantly affected by repeat stress (*H* = 5.6, *P* = 0.0608, d.f. = 2; Fig. S5B). Social status did not affect brain *gr* mRNA levels (F_2, 18_ = 2.871, *P* = 0.0828; Fig. 5C).

**Figure 5.**
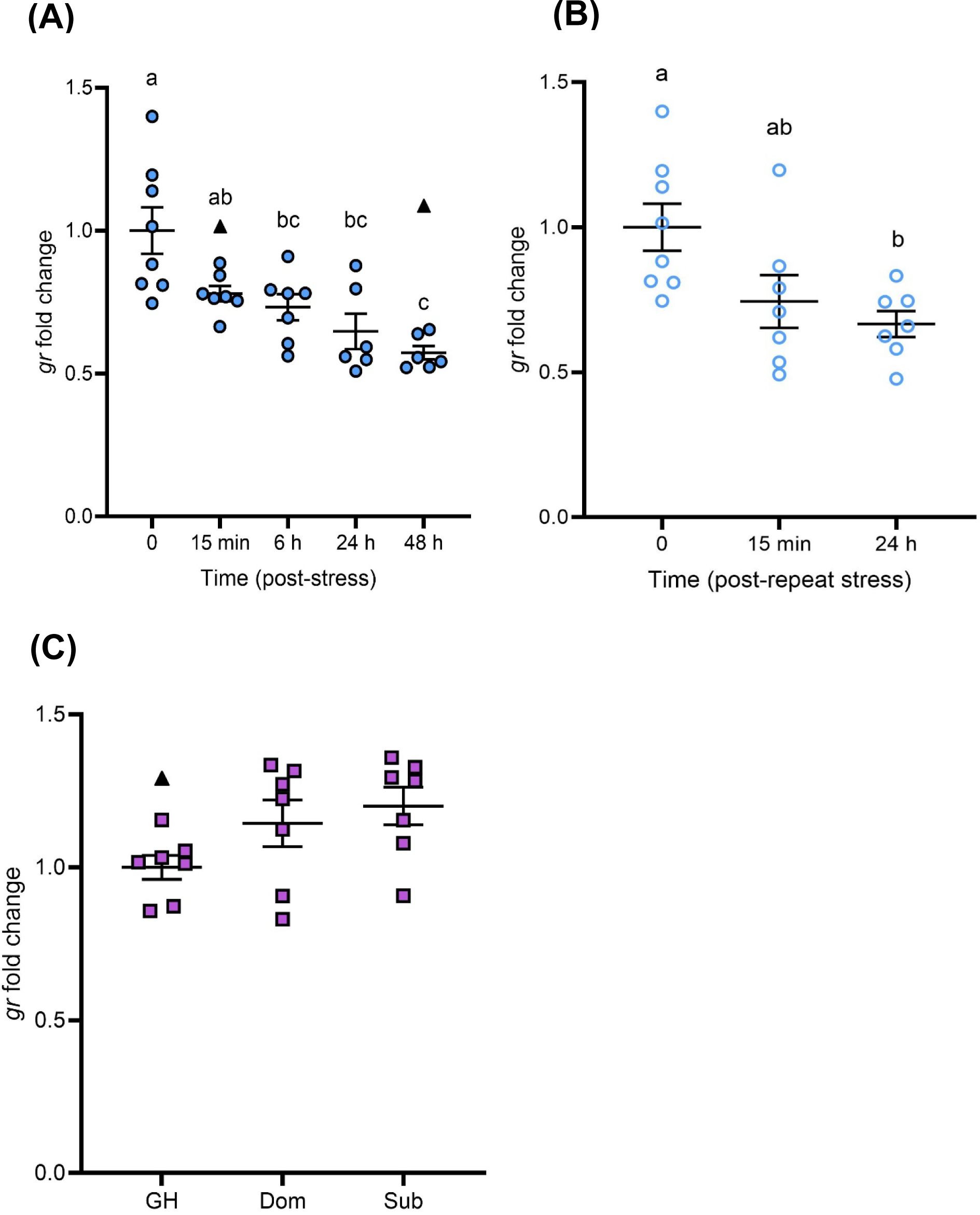
Effects of stress exposure on *gr* transcript abundance in the adult zebrafish brain. **(A)** Zebrafish were exposed to air for 1-min and allowed to recover for up to 48 h (closed blue circles). **(B)** For a separate group of fish, the stressor was repeated 24 h after the first stressor and fish were allowed to recover for 15 min or 24 h (open blue circles). **(C)** Dominant (Dom) and subordinate (Sub) zebrafish after 24 h of social stress; group-housed (GH) fish are included as a control (purple squares). Datapoints identified as statistical outliers were removed prior to analysis, and are shown as black triangles. Gene transcript abundance was normalized to the mean expression of two housekeeping genes (*ef1α* and *rpl8*). Data is shown as individual data points with a solid line and whiskers representing the mean ± SEM (*gr* transcript abundance, acute stress n = 6-8; repeat stress n = 7-8; social stress n = 7). Changes in *gr* transcript abundance following stress exposure were determined using a one-way ANOVA with a Tukey’s *post-hoc* test (single and repeat stress *P* < 0.05; social stress *P* > 0.05). Letters denote significant differences among groups within a panel; groups which have the same letter are not significantly different from one another.

Brain *mr* mRNA abundance was significantly different among timepoints following acute stress exposure (F_4, 28_ = 3.513, *P* = 0.0191; Fig. 6A). After 6 h recovery, *mr* mRNA levels were 1.7-fold lower than pre-stress levels (*P* = 0.0137) but recovered to baseline by 24 h. In contrast, *mr* mRNA abundance within the pooled telencephalons was not significantly affected by exposure to an acute stressor (*H* = 2.081, *P* = 0.5558, d.f. = 3; Fig, S5C). Repeat stress significantly affected brain *mr* mRNA abundance (F_2,18_ = 8.521, *P* = 0.0025; Fig. 6B), where a significant 1.6-fold reduction in *mr* RNA levels occurred at 15 min and 24 h recovery, relative to baseline (*P* = 0.0067). Telencephalon *mr* mRNA abundance was not significantly affected by repeat stress exposure (F_2,6_ = 2.611, *P* = 0.1530; Fig. S5D). Social status did not affect brain *mr* mRNA levels (F_2, 19_ = 0.7530, *P* = 0.4850; Fig. 6C).

**Figure 6.**
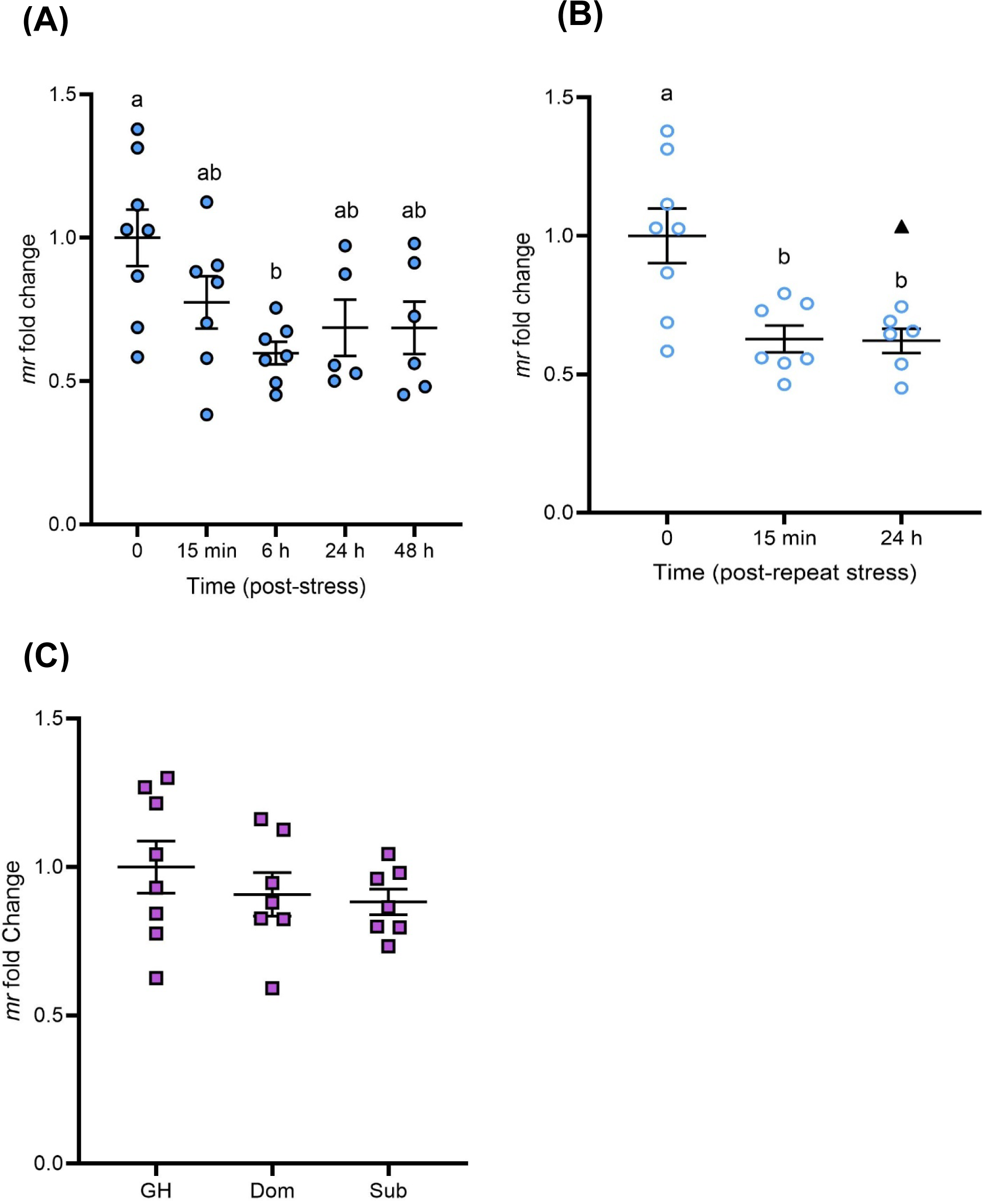
Effects of stress exposure on *mr* transcript abundance in the adult zebrafish brain. **(A)** Zebrafish were exposed to air for 1-min and allowed to recover for up to 48 h (closed blue circles). **(B)** For a separate group of fish, the stressor was repeated 24 h after the first stressor and fish were allowed to recover for 15 min or 24 h (open blue circles). **(C)** Dominant (Dom) and subordinate (Sub) zebrafish after 24 h of social stress; group-housed (GH) fish are included as a control (purple squares). Datapoints identified as statistical outliers were removed prior to analysis, and are shown as black triangles. Gene transcript abundance was normalized to the mean expression of two housekeeping genes (*ef1α* and *rpl8*). Data is shown as individual data points with a solid line and whiskers representing the mean ± SEM (*mr* transcript abundance, single stressor n = 5-8; repeat stressor n = 6-8; social stress n = 7-8). Changes in *mr* transcript abundance following stress exposure were determined using a one-way ANOVA with a Tukey’s *post-hoc* test (single and repeat stress *P* < 0.05; social stress *P* > 0.05). Letters denote significant differences among groups within a panel; groups which have the same letter are not significantly different from one another.

## Discussion

This study demonstrates the dynamic and stressor-specific regulation of Hsd11b2 expression in the zebrafish brain. An acute increase in cortisol generated a sustained up-regulation of brain Hsd11b2 expression at the transcript and protein level for more than 24 h after the stress. Under the same time frame, a repeated acute stressor blunted this response while a chronic increase in cortisol failed to elicit any changes in brain Hsd11b2 expression. Importantly, the changes in Hsd11b2 elicited by acute stress coincided with reduced *gr* and *mr* expression as well as increased brain cell proliferation. These results suggest that intracellular cortisol signaling capacity can be tempered through a combination of increased cortisol inactivation and receptor down-regulation. In turn, these changes would temporarily spare the inhibitory effects of stress on neurogenesis and may even potentiate pro-neurogenic signaling. Thus, this study supports Hsd11b2 as an agent for mediating stress-specific changes in brain cell proliferation.

### Stressor-specific regulation of hsd11b2 expression in the adult zebrafish brain

Acute and chronic stressors induce characteristic but distinct changes in circulating glucocorticoids (Madliger and Love 2014; Romero and Gormally, 2019). In zebrafish, a substantial increase in cortisol is observed within minutes of a fish experiencing an acute stressor and then returns to basal levels within hours of recovery (Ramsay et al., 2009; Fuzzen et al., 2010; Alderman and Vijayan, 2012). This is consistent with results of the present study, where plasma cortisol was significantly elevated 15 min after a brief air exposure and fully recovered by 24 h. This cortisol profile was exactly replicated when the stressor was repeated one day later; hence designating these regimes as acute and repeat acute stressors, respectively. In contrast, chronic stressors that persist for hours or days are frequently associated with sustained increases in circulating cortisol that last at least until the stressor abates. Dyadic fish pairings can serve as a chronic stressor in laboratory studies owing to the rapid establishment of distinct, robust behavioural phenotypes and elevated cortisol in socially subordinate fish throughout the interaction (Øverli et al., 1999; Sloman et al., 2001; Alderman et al., 2008; Bernier et al., 2008; Filby et al., 2010; Tea et al., 2019; but see Höglund et al., 2001; Pavlidis et al., 2011; Sørensen et al., 2012; Jeffrey and Gilmour 2016). In the present study, cortisol levels were higher in subordinate relative to dominant male zebrafish after 24 h of continuous interaction, hence this regime was designated a chronic stressor.

The distinct cortisol profiles under acute and chronic stress scenarios are thought to underlie the biphasic outcomes of elevated cortisol (Herman, 2013). For example, whereas acute stressors stimulate energy mobilization (Mommsen et al., 1999; Schreck and Tort, 2016), chronic stressors can result in prolonged catabolic responses that lead to growth inhibition (Sadoul and Vijayan, 2016). While this framework applies to many known actions of cortisol, including neurogenesis (Sørensen et al., 2013; Saaltink and Vreugdenhil, 2014), the cellular mechanisms that underpin such biphasic effects are not fully understood. Cortisol catabolism by Hsd11b2, which is irreversible in fish (Tsachaki et al., 2017), may play a role. At the organismal level, the absence of Hsd11b2 activity induced by pharmacological inhibition (Alderman and Vijayan, 2012) or CRISPR/Cas9-mediated knockout (Theodoridi et al., 2021) increased basal cortisol levels and delayed cortisol recovery following acute stress, highlighting a key role for Hsd11b2 in the physiological response to stress.

Stress-induced changes in *hsd11b2* expression in zebrafish have been observed in several cortisol target tissues including the head kidney (Fuzzen et al., 2010), gill and ovary (Lim and Bernier, 2022), as well as the brain (Alderman and Vijayan, 2012; Madaro et al., 2016; Huang et al., 2019). Nevertheless, there is considerable variation in the nature of this response, no doubt influenced by within-study restrictions on tissue/temporal sampling and across-study differences in stressor application. For example, Fuzzen et al. (2010) tracked *hsd11b2* mRNA levels in the head kidney of zebrafish during a 1 h agitation stress and 20 min recovery and showed that the transient increase in gene expression closely aligned with the cortisol profile in these fish; however, these temporal changes were reported within the context of a single stressor and tissue. Lim and Bernier (2022) demonstrated stressor- and tissue-specific changes in *hsd11b2* mRNA abundance in the ovaries and gills of zebrafish; however, measurements were taken at a single timepoint following chronic exposure to cycling hypoxia, temperature, or combination stressors. The present study advances our understanding of the dynamic changes in Hsd11b2 expression associated with stressor-specific cortisol profiles and includes responses at both mRNA and protein levels. Specifically, we show that recovery from an acute stressor was marked by a persistent up-regulation of Hsd11b2 gene and protein expression in the brain through at least 48 h, supporting the previously reported elevation in Hsd11b2 enzymatic activity in zebrafish brains 24 h after experiencing a single air exposure (Alderman and Vijayan, 2012). Intriguingly, this up-regulation in Hsd11b2 expression was quickly suppressed in fish that experienced a second air exposure stressor, as both *hsd11b2* mRNA and protein abundance returned to baseline levels after 24 h recovery from repeat stress, the temporal equivalent of 48 h recovery in the single acute stress experiment. Indeed, the apparent increase in Hsd11b2 protein abundance at 15 min post repeat stress may simply be a relic of the initial air exposure (temporal equivalent of 24 h recovery from acute stress). In contrast, the chronic stress of social subordination imposed a modest increase in brain *hsd11b2* expression at the gene but not protein level, aligning with previous work in rainbow trout after 96 h of social subordination (Best et al., 2023). Taken together, these results support rapid, experience-based changes in the capacity of the brain to catabolize intracellular cortisol, and subsequently, regulate downstream effects of cortisol signaling.

### A mechanism for regulating the effects of stress on adult neurogenesis

Hsd11b2 is poised to contribute to context-specific regulation of adult neurogenesis by cortisol in zebrafish, given its localization to neurogenic niches (Alderman and Vijayan, 2012) and the stressor-induced changes in its expression (present study; Alderman and Vijayan, 2012; Best et al., 2023). Here, we provide novel support for this role by combining evidence of temporal changes in cortisol and Hsd11b2 expression with measures of brain cell proliferation. Consistent with the purported pro-neurogenic effects of acute stress in mammals (Thomas et al., 2006; Dagyte et al., 2009; Kirby et al., 2013; So et al., 2017) and teleost fish (von Krogh et al., 2010 Johansen et al., 2011; Zhang et al., 2020), we show that the expression of *pcna* in the telencephalon and remaining brain regions increases within 15 min of recovery from acute stress, concomitant with elevated cortisol. There were also more BrdU+ cells in the telencephalon 1.5 h after air exposure, confirming a link between *pcna* expression and mitosis. Notably, the increase in brain Hsd11b2 protein expression (present study) and activity (Alderman and Vijayan, 2012) that was also induced by acute stress was delayed relative to the cortisol response, suggesting that the early aftermath of stress is marked by a limited capacity to buffer intracellular cortisol levels in the brain. Given that repeat exposure to the acute stressor did not further promote brain cell proliferation, we propose that the increased capacity for cortisol catabolism realized by elevated Hsd11b2 prevented a second pro-neurogenic response in the repeat stress group.

Under chronic stress, brain *pcna* expression was unchanged in fish following 24 h of social subordination. This result was unexpected, given that decreased *pcna* expression was observed in the hypothalamus, optic tectum, and telencephalon of rainbow trout following 72 h of social stress (Johansen et al., 2012). Similarly, fewer BrdU+ cells were observed in the telencephalons of subordinate male zebrafish that were exposed to the mitotic marker after 24 h of subordination and euthanized one day later (Tea et al., 2019). In mammals, *pcna* expression has been shown to be highly transient during the cell cycle, with increased expression during S-phase followed by an immediate decline in M-phase (Ino and Chiba, 2000). In our acute stress experiment, we also found that *pcna* expression declined within hours of the initial increase. Thus, single timepoint assessment of subordinate fish may be insufficient to capture a change in *pcna* expression. Nevertheless, the absence of an Hsd11b2 response under chronic stress, as reported here, is consistent with our hypothesis. With a limited or status quo capacity to catabolize cortisol, intracellular cortisol in NSPCs could reach or exceed the threshold required to induce anti-neurogenic signaling.

Variation in receptor abundance is another mechanism by which context-specific responses to cortisol can be realized in target tissues. At the same time, Gr and Mr have different ligand binding affinities in fish (Bury et al., 2003; Greenwood et al., 2003; Sturm et al., 2005), as is seen in mammals (Koning et al., 2019). Thus, while both Gr and Mr are widely expressed in the fish brain (Wendelaar Bonga, 2011), the relative proportion of high-affinity Mr to low-affinity Gr may also influence the nature of cortisol signaling in target cells. Indeed, stress is known to impart brain region-specific changes in *gr* and *mr* mRNA abundance (Johansen et al., 2011; Alderman and Vijayan, 2012; Pavlidis et al., 2015; Madaro et al., 2016; Molteson et al., 2016; Kiilerich et al., 2018; Best et al., 2023). In rainbow trout selected for distinct stress coping styles, fish characterized by a strong cortisol response to acute stress had lower *mr* but similar *gr1* and *gr2* expression across several brain regions compared to fish characterized by a low cortisol response to acute stress (Johansen et al., 2011). These differences in receptor expression are thought to explain, at least in part, the distinct behavioural and physiological phenotypes of these fish (Johansen et al., 2011), including stress-induced changes to *pcna* expression in the telencephalon (Johansen et al., 2012). In the present study, there was a consistent downregulation of brain *gr* and *mr* expression after both acute and repeat acute stress exposure, but no changes under chronic stress. Thus, although the two cortisol receptors did not show differential responses, these results support receptor abundance as a mechanism for stressor-specific effects of cortisol in the brain. Interestingly, the timing of receptor expression changes was delayed relative to the initial cortisol response following air exposure, as occurred with Hsd11b2; however, *gr* and *mr* downregulation persisted through repeat stress whereas Hsd11b2 recovered. Taken together, these results emphasize that the capacity of NSPCs for cortisol catabolism and receptor-mediated signaling is influenced by experience.

## Conclusion

Stress imparts myriad effects on animal physiology, including in the brain. Here, the influence of cortisol on the cell cycle can alter neural circuits that are populated by new neurons derived from adult NSPCs, providing a mechanism for experience-based learning. In animals with a high regenerative capacity, like the zebrafish, regulating the influence of cortisol on this process may be necessary to strike a balance between continuity and plasticity in the neural circuitry of the brain. This study presents evidence that Hsd11b2 is differentially regulated in the zebrafish brain under acute and chronic stress. We propose the Hsd11b2 response as a novel mechanism through which NSPCs may modulate rates of cortisol catabolism and subsequently, context-specific changes to cell proliferation in the post-natal brain. While we provide clear evidence that Hsd11b2 is regulated by stress, it remains unknown whether the changes reported in this study are themselves mediated by increased cortisol or by alternate pathways. In support of the former, putative glucocorticoid response elements have been identified in the promotor regions of both mammalian and zebrafish *hsd11b2* genes (Surjit et al., 2011; Alderman and Vijayan, 2012). However, future work could also explore other molecular and cellular signaling pathways such as DNA methylation, microRNAs, and the p38 mitogen-activated protein kinase (P38 MAPK) pathway. These pathways have been implicated as potential mechanisms for the regulation of HSD11B2 in mammals (Sharma et al., 2009; Peña et al., 2012; Razaei et al., 2014) and warrant further exploration in fish. Studies that increase our understanding of the stress-specific pathways controlling the regulation of Hsd11b2 will provide further insight into how zebrafish utilize endogenous regulators to directly mediate changes to adult neurogenesis and subsequently, help elucidate how neuroplasticity is maintained throughout their life.

## Acknowledgements

We thank Dr. Kathleen M. Gilmour (University of Ottawa) who generously provided the custom Hsd11b2 antibody. We thank Matt Cornish and Mike Davies (Hagen Aqualab) for their assistance with experimental set-ups. In addition, we thank Chengcheng Zhang, Julia Bourdeau, Amanda Wiseman, Elizabeth Manchester, Jared Shaftoe, and Reece Long for their assistance with animal care.

## Data Availability

Raw data is available on request.

## List of Abbreviations

ANOVA: Analysis of variance
AU: Arbitrary unit
BrdU: 5’-bromo-2’-deoxyuridine
cDNA: Complementary deoxyribonucleic acid
Ct: Cycle threshold
Ef1α: Elongation factor 1α
ELISA: Enzyme-linked immunosorbent assay
Gr: Glucocorticoid receptor
Hsd11b2: 11β-hydroxysteroid dehydrogenase type 2
Hsd20b2: 20β-hydroxysteroid dehydrogenase type 2
IQR: Interquartile Range
Mr: Mineralocorticoid receptor
mRNA: Messenger ribonucleic acid
MS-222: Tricaine methane sulfonate
_NaHCO3_: Sodium bicarbonate
Non-RT: Non-reversed transcribed
NSPC: Neural stem and progenitor cells
PBS: Phosphate-buffered saline
PBST: PBS containing 0.05% Triton X-100
PC1: First principal component
PCA: Principal components analysis
Pcna: Proliferating cell nuclear antigen
Rpl8: Ribosomal protein L8
RT-qPCR: Reverse transcription quantitative real-time polymerase chain reaction
SDS-PAGE: Sodium dodecyl sulfate-polyacrylamide gel electrophoresis
TBS: Tris-buffered saline
TBST: Tris-buffered saline containing 0.05% Triton X-100

## Supplemental Information

**Supplemental Figure S1.**
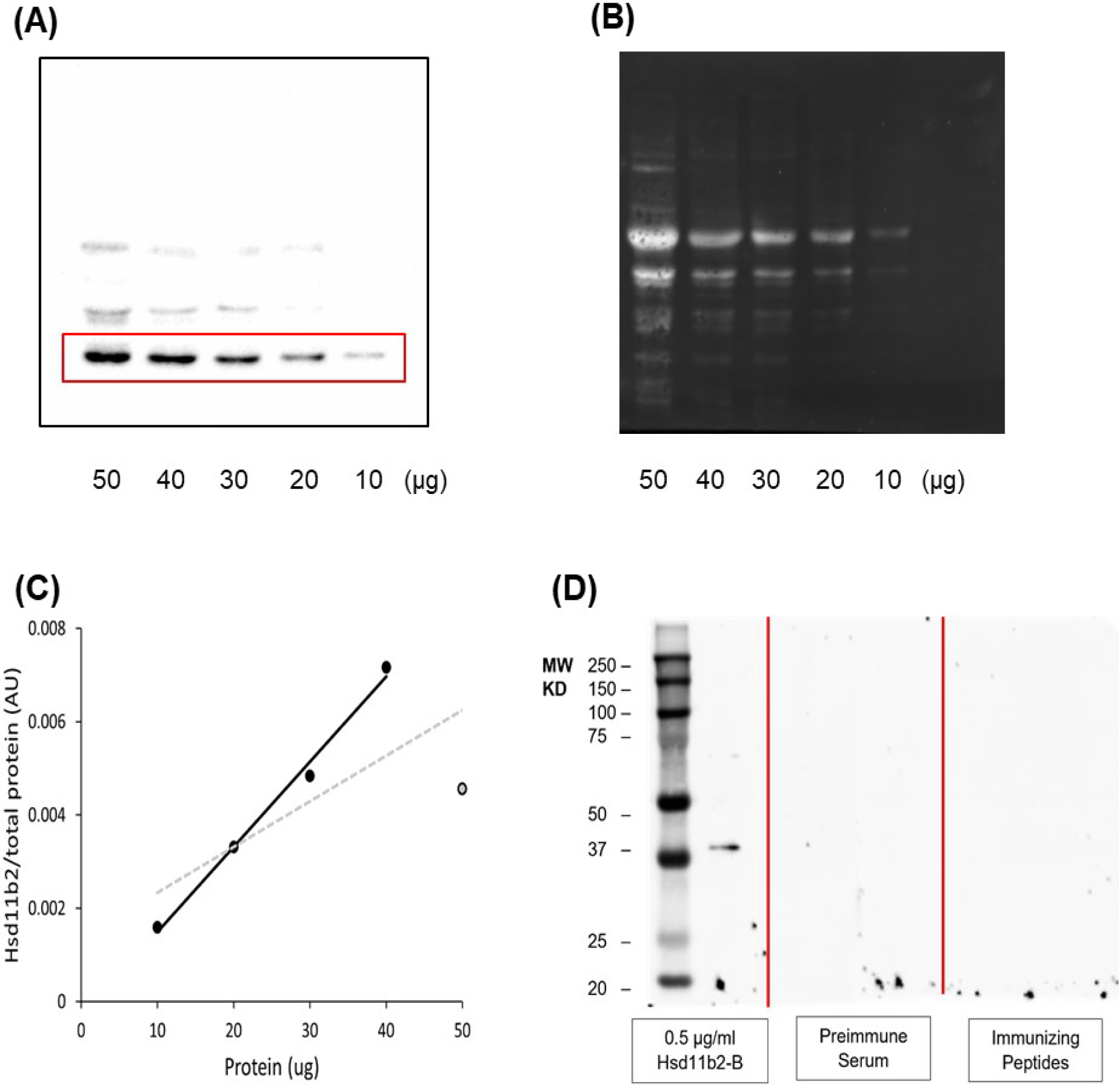
**(A)** A western blot where increasing amounts of zebrafish brain homogenate (10 - 50 µg) were loaded on 10%SDS-PAGE gel and transferred to a nitrocellulose membrane. ECL detection of Hsd11b2 (bands outlined by red rectangle) using the chicken anti-Hsd11b2 antibody. **(B)** Fluorescent detection of total protein on the same blot using SYPRO Ruby Red Protein Blot Stain. **(C)** A linear regression analysis of Hsd11b2/total protein band intensity in proportion to increased protein loading showing the linear range of detection, represented by the black trendline (10 - 40 µg, R^2^ = 0.9913), and that above 40 µg, the signal begins to plateau and sample loading does not exhibit the expected increase in band intensity, represented by the grey dashed trendline (10 - 50 µg, R^2^ = 0.5673). **(D)** A western blot where 20 µg aliquots of zebrafish brain homogenate were loaded into each lane and then divided into 3 pieces following transfer (red line). From left to right, the split blots were incubated in 0.5 µg ml^-1^ Hsd11b2-B (38-KDa), pre-immune serum, and 0.5 µg mL^-1^ Hsd11b2 containing 5 µg mL^-1^ immunizing peptide. There was no immunoreactivity observed in the pre-immune or immunizing peptide sections proving the specificity of the Hsd11b2-B antibody.

**Supplemental Figure S2.**
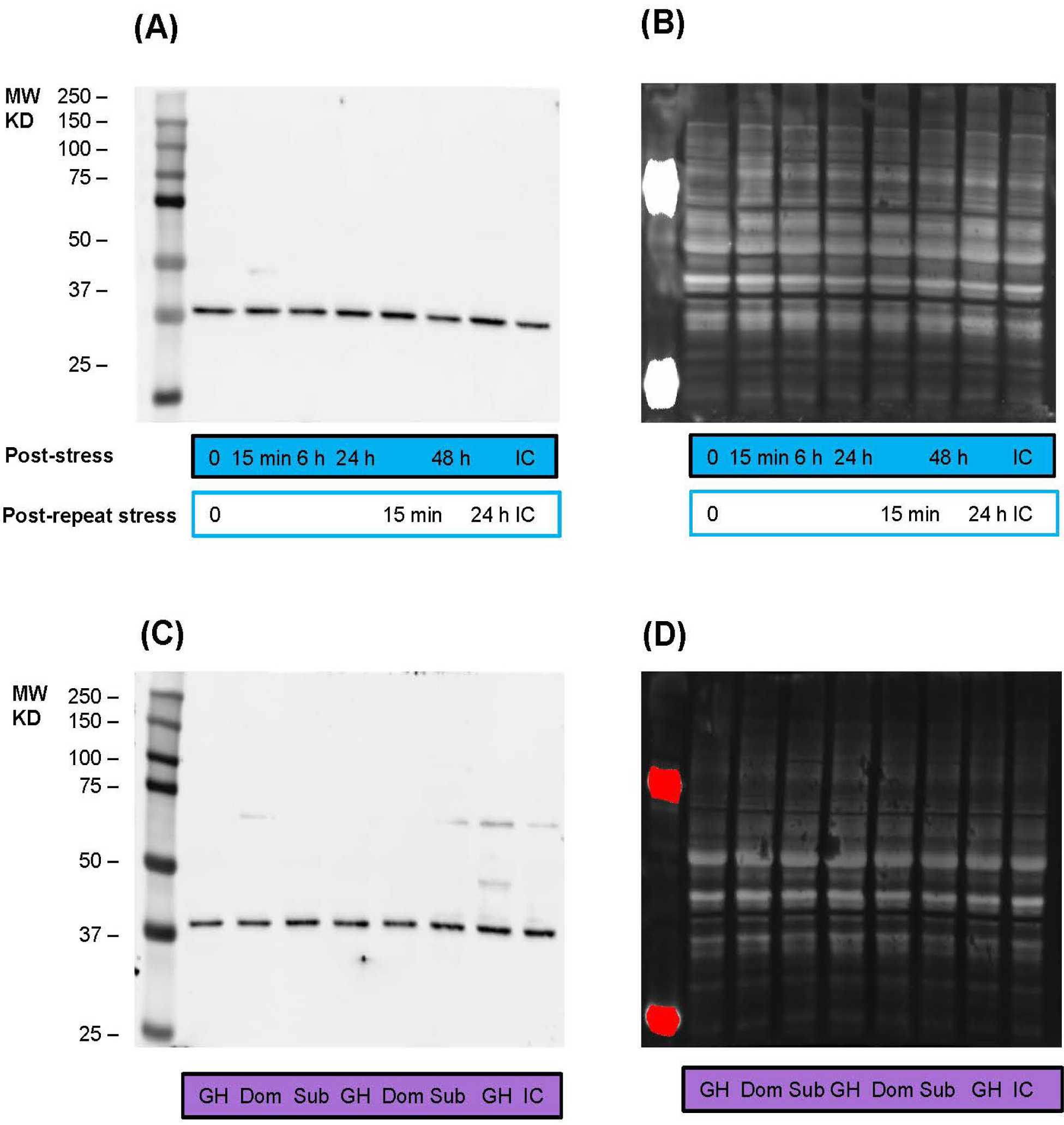
Representative western blots showing **(A)** Hsd11b2 protein expression determined using Hsd11b2-B antibody, and **(B)** total protein determined using SYPRO Ruby Red Protein Blot Stain following exposure to a single (solid blue) and repeat (blue outline) 1-min air exposure stressor and recovery in the brains of adult zebrafish. Representative western blots of **(C)** Hsd11b2 protein expression determined using Hsd11b2-B antibody, and **(D)** total protein determined using SYPRO Ruby Red Protein Blot Stain following 24 h of social stress in the brains of dominant (Dom) and subordinate (Sub) zebrafish; group-housed (GH) fish are included as a control. IC = internal control from pooled brain homogenates.

**Supplemental Figure S3.**
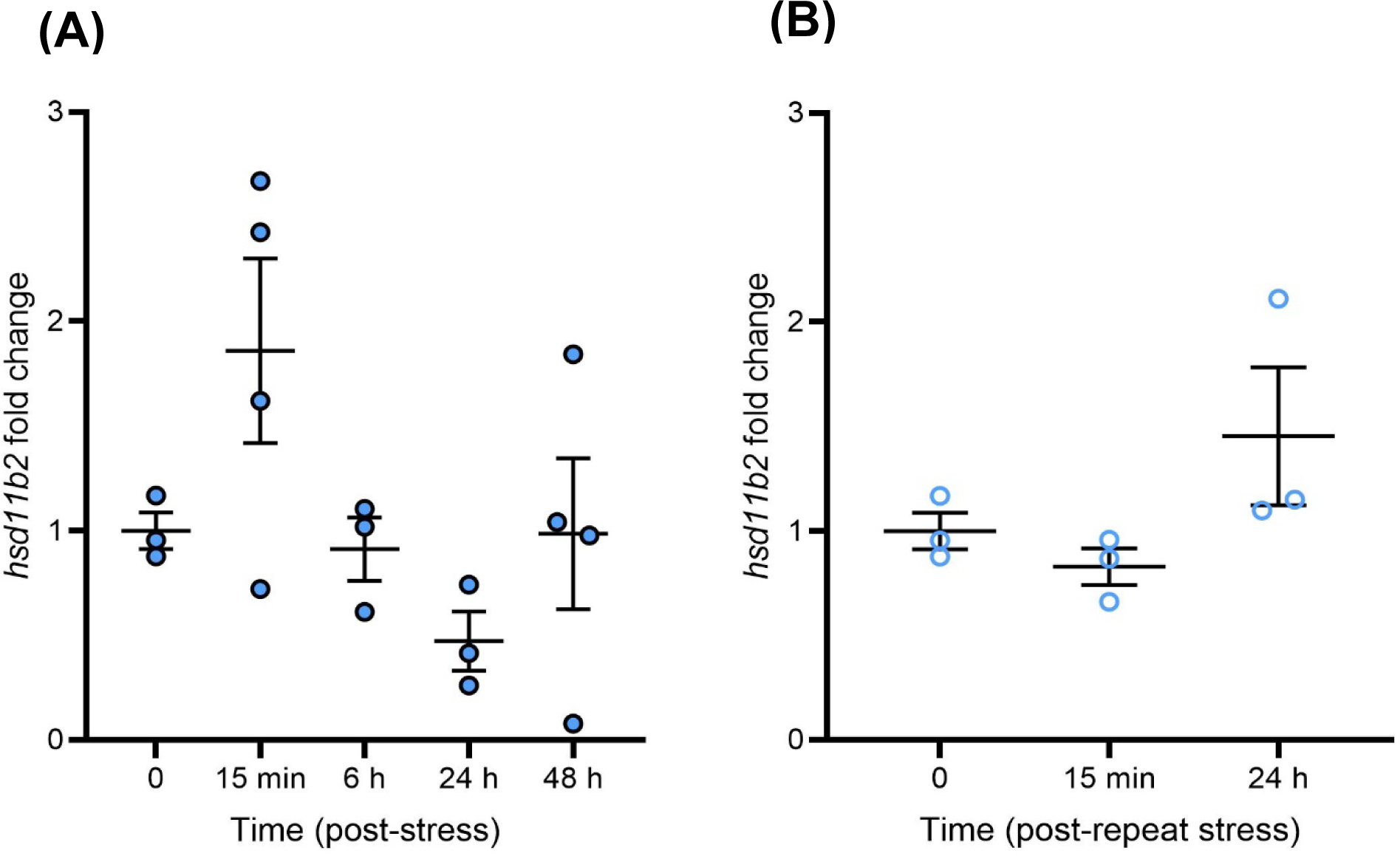
Effects of an acute (closed blue circles) and repeat (open blue circles) 1-min air exposure stressor on *hsd11b2* transcript abundance in the zebrafish telencephalon (n=2 fish pooled per sample). **(A)** Pooled telencephalons *hsd11b2* transcript abundance after exposure to an acute, and **(B)** repeat stressor. Gene transcript abundance is normalized to the mean expression of two housekeeping genes (*ef1α* and *rpl8*). Data is presented as individual data points, with solid lines and whiskers representing the mean ± SEM for each timepoint (*hsd11b2* transcript abundance acute stressor n = 3-4, repeat stressor n = 3). Differences in *hsd11b2* mRNA levels following acute and repeat stress exposure were determined using a one-way ANOVA with a Tukey’s *post-hoc* test (*P* > 0.05).

**Supplemental Figure S4.**
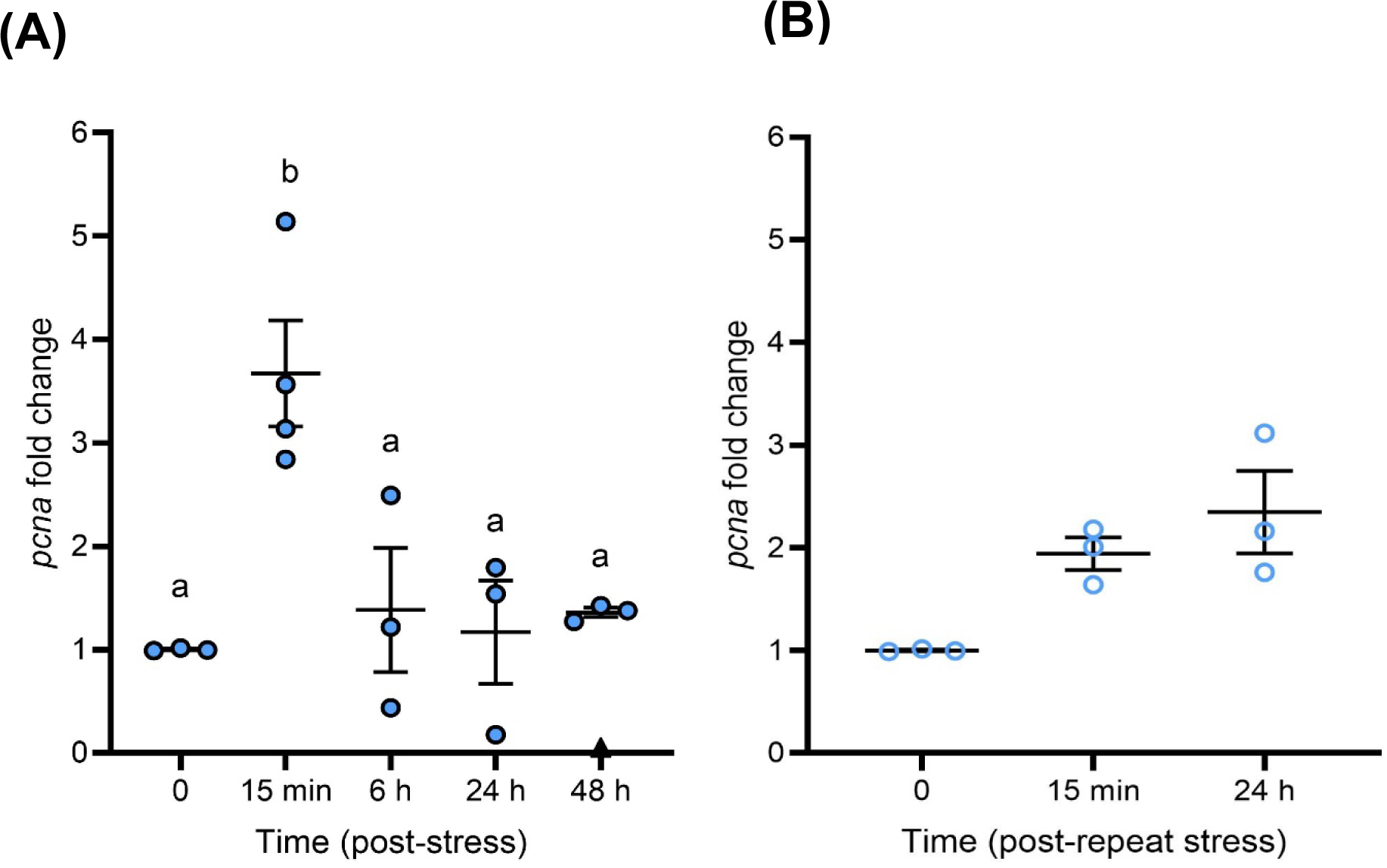
Effects of an acute (closed blue circles) and repeat (open blue circles) 1-min air exposure stressor on *pcna* transcript abundance in the pooled zebrafish telencephalon. **(A)** Pooled telencephalons *pcna* transcript abundance after exposure to an acute, and **(B)** repeat stressor. Gene transcript abundance is normalized to the mean expression of two housekeeping genes (*ef1α* and *rpl8*). Data is presented as individual data points, with solid lines and whiskers representing the mean ± SEM for each timepoint (*pcna* transcript abundance acute stressor n = 3-4, repeat stressor n = 3). Outliers removed from treatment groups represented by black triangles. Differences in *pcna* mRNA levels following acute (*P* < 0.05) and repeat stress (*P >* 0.05) exposure were determined using a Kruskal Wallis test followed by Dunn’s *post-hoc* test. Letters indicate significant differences among groups within a panel; groups which have the same letter are not significantly different from one another.

**Supplemental Figure S5.**
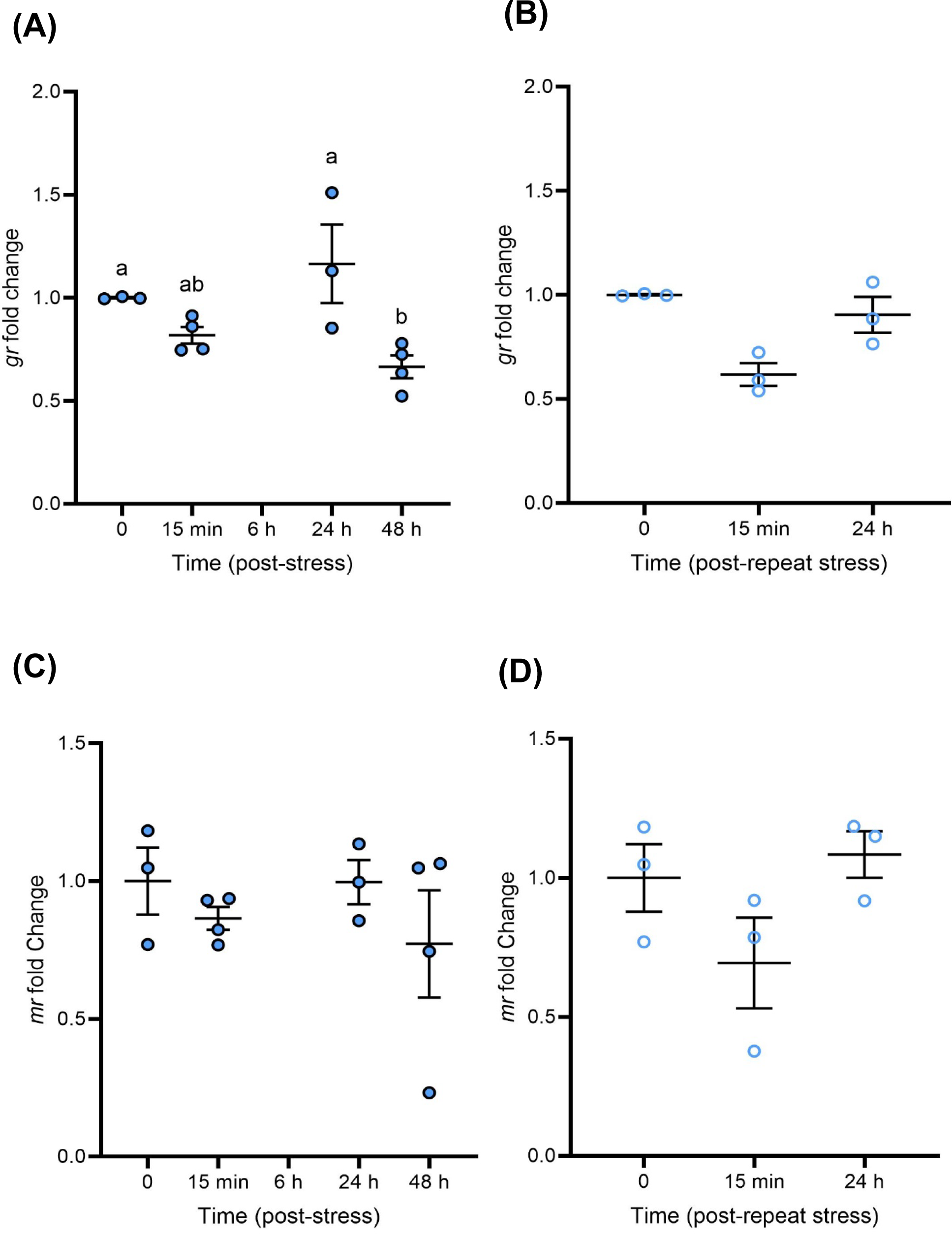
Effects of an acute (closed blue circles) and repeat (open blue circles) 1-min air exposure stressor on *gr* and *mr* transcript abundance in the pooled zebrafish telencephalon. **(A)** Pooled telencephalons *gr* transcript abundance after exposure to an acute, and **(B)** repeat stressor. **(C)** Pooled telencephalons *mr* transcript abundance after exposure to an acute, and **(D)** repeat stressor. Gene transcript abundance is normalized to the mean expression of two housekeeping genes (*ef1α* and *rpl8*). Data is presented as individual data points, with solid lines and whiskers representing the mean ± SEM for each timepoint (*gr* transcript abundance acute stressor n = 2-4, repeat stressor n = 3; *mr* transcript abundance acute stressor n = 2-4, repeat stressor n = 3). The 6 h post-stress recovery timepoint is omitted for *gr* and *mr* mRNA abundance due to an insufficient sample size. Differences in *gr* (acute stress *P* < 0.05, repeat stress *P >* 0.05) and *mr* (P > 0.05) mRNA levels following acute and repeat stress exposure were determined using a either a one-way ANOVA followed by Tukey’s *post-hoc* test or a Kruskal Wallis test followed by Dunn’s *post-hoc* test. Letters indicate significant differences among groups within a panel; groups which have the same letter are not significantly different from one another.

**Supplemental Figure S6.**
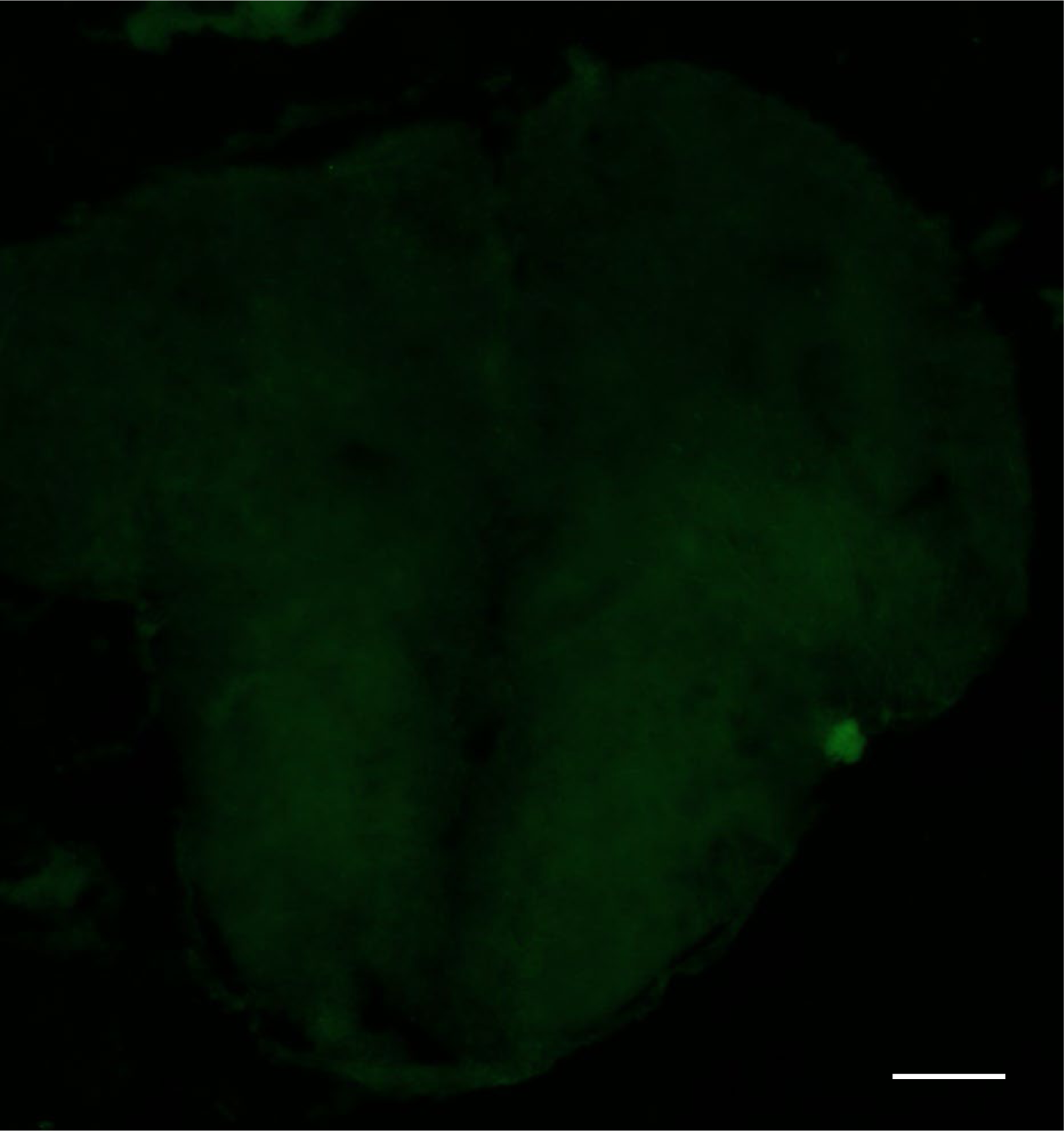
Representative image of a cross-section through a zebrafish telencephalon acting as a no primary control. This representative image coincides with the cross-sectional view of the rostral telencephalon Level 98 (Wullimann et al., 1996). There was no immunoreactivity observed proving the specificity of the the primary antibody, mouse anti-BrdU (Developmental Studies Hybridoma Bank). Scale bar = 200 µm.

